# Validation of Data Acquisition and Phase Estimation for Quantitative Susceptibility Mapping with a Rotating-Tube Phantom

**DOI:** 10.1101/2021.06.29.450224

**Authors:** Kathryn E. Keenan, Ben P. Berman, Slavka Carnicka, Stephen E. Russek, Wen-Tung Wang, John A. Butman, Dzung L. Pham, Joseph Dagher

**Author notes:** NIST required disclaimer: Certain commercial instruments and software are identified to specify the experimental study adequately. This does not imply endorsement by NIST or that the instruments and software are the best available for the purpose.

## Abstract

**Purpose:** Quantitative Susceptibility Mapping (QSM) is an MRI tool with the potential to reveal pathological changes from magnetic susceptibility measurements. Before phase data can be used to recover susceptibility (Δ*χ*), the QSM process begins with two steps: data acquisition and phase estimation. We assess the performance of these steps, when applied without user intervention, on several variations of a phantom imaging task.

**Approach:** We used a rotating-tube phantom with five tubes ranging from Δ*χ*=0.05 ppm to Δ*χ*=0.336 ppm. MRI data was acquired at nine angles of rotation for four different pulse sequences. The images were processed by 10 phase estimation algorithms including Laplacian, region-growing, branch-cut, temporal unwrapping and maximum-likelihood methods. We analyzed errors between measured and expected phase using the probability mass function and Cumulative Distribution Function.

**Results:** Repeatable acquisition and estimation methods were identified based on the probability of relative phase errors. For single-echo GRE and segmented EPI sequences, a region-growing method was most reliable with Pr(relative error<0.1)=0.95 and 0.90 respectively. For multi-echo sequences, a Maximum-Likelihood method was most reliable with Pr(relative error<0.1)=0.97. The most repeatable multi-echo methods outperformed the most repeatable single-echo methods.

**Conclusions:** We found a wide range of repeatability and reproducibility for off-the-shelf MRI acquisition and phase estimation approaches. The error was dominated in many cases by spatially discontinuous phase unwrapping errors. Any post-processing applied on erroneous phase estimates, such as QSM’s background field removal and dipole inversion, would suffer from error propagation. Our paradigm identifies methods that yield consistent and accurate phase estimates that would ultimately yield consistent and accurate Δ*χ* estimates.

## 1 Introduction

Quantitative Susceptibility Mapping (QSM)^1^ is a method to estimate magnetic susceptibility of tissue from the phase of the magnetic resonance (MR) signal. QSM has potential clinical utility for characterizing neurological diseases^2-4^, blood oxygen content^5^, and iron overload in the heart and liver^6^.

Repeatability and reproducibility of QSM has been assessed in phantoms and human subjects using different scanners, magnetic field strengths, and data processing methods. While some studies report high repeatability^7-14^, both *in vivo* and in phantoms, recent *in vivo* studies report lower reproducibility across MRI scanners with the same data processing method^15^ and across QSM algorithms using the same input data^16^. These conflicting results limit the clinical adoption of QSM.

A typical QSM process requires four steps: data acquisition (Step 1), phase estimation (Step 2), background field removal (Step 3) and magnetic susceptibility reconstruction (Step 4). Recently proposed methods combine Steps 2-4 into fewer steps^17^.

In its standardization efforts, the QSM community has actively evaluated competing methods^13, 18, 19^, in particular for methods in Steps 3 and 4 of the process^16^. However, the selection of the “best” QSM method is difficult for various reasons: a) the appropriate definition of a quality metric, e.g. accuracy vs repeatability; b) competing quality metrics that favor different algorithms^16^; c) the lack of a gold standard *in vivo*; d) algorithm performance depends on imaging application (*in vivo* vs phantoms); and e) the large number of methods for each QSM step, which would render any exhaustive validation effort to be combinatorial and quickly untenable.

We present an experimental setup that allows for an exhaustive quantitative analysis of all four QSM steps. This framework uses a rotating-tube phantom design introduced in Erdevig et al^20^, which uses tubes that rotate, within a background solution, relative to the main magnetic field, B_0._ The design enables the analysis of MRI data obtained in objects at any orientation, using common QSM techniques. The closed-form theoretical relationship between the magnetic field and magnetic susceptibility in the sample allows for mapping the magnetic field to magnetic susceptibility without having to solve the dipole inversion problem^21^.

Our contributions include a) a framework for evaluation of repeatability and reproducibility of QSM algorithms; and b) rigorous analysis of common methods in Steps 1 & 2 of the QSM process. We focus on the performance of QSM Steps 1 & 2 for the following reasons:

a. There exist a large number of methods in each of the four QSM steps. We replicate here approximately 90 different combinations of Steps 1 & 2 methods. If we analyze all four steps simultaneously, the resulting data set would grow combinatorically and become difficult to interpret.
b. Errors introduced early in the QSM process propagate downstream and have been overlooked in validation studies. For example, Olsson et al used only one acquisition sequence and one phase estimation algorithm^18^.
c. It is well understood that phase unwrapping, a common algorithm used in Step 2, is a non-deterministic polynomial-time (NP)-hard problem (in two dimensions and higher) that often relies on user intervention and careful parameter tuning. Therefore, it is important to isolate the impact of such a problem in the QSM processing methods.
d. Susceptibility weighted imaging^22, 23^, electrical properties tomography^24^, thermometry^25^, flow^26^, and elastography^27^ use MR phase information, and this work can inform those applications.

We analyze Steps 1 and 2 of the QSM process to understand which are sufficiently robust to be executable without user intervention, and independent of scanner, sequence and other parameter variations.

## 2 Methods

We used a rotating-tube phantom (Fig. 1a) to explore the reproducibility of phase estimates obtained after Steps 1 & 2. MRI data was acquired with different pulse sequences (Step 1), at nine angles, and the performance of a wide variety of phase estimation methods was compared to theory. The rotating-tube phantom was designed to take advantage of the analytical model for a long cylinder at an angle, θ, with respect to B_0_. The internal z-axis field offset can be derived from Maxwell’s equations and is shown to be^21^:

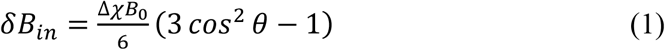

where Δ*χ* is the susceptibility difference between the inside and outside of the cylinder, and *χ*^2^ terms are ignored.

**Fig. 1.**
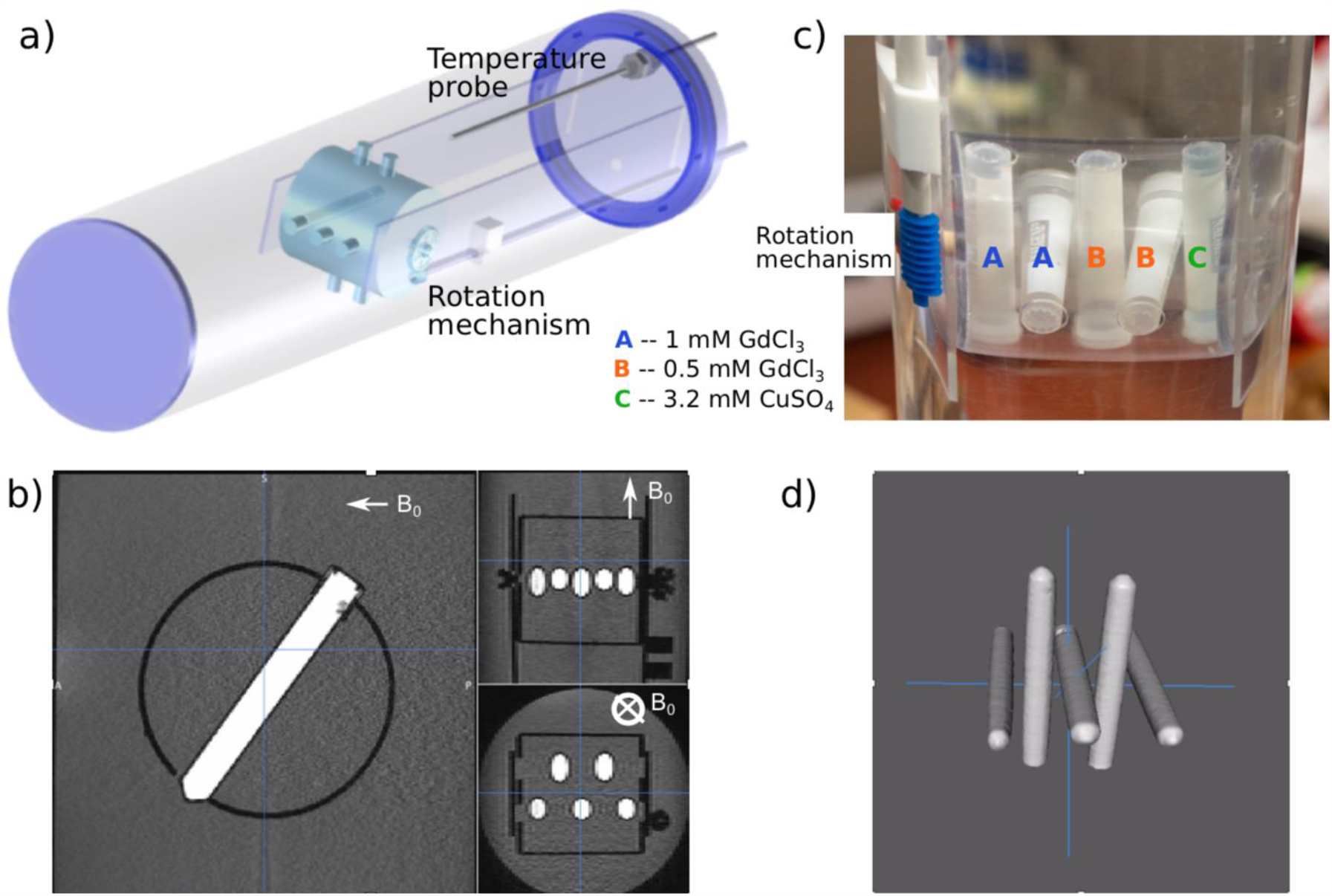
(a) 3D diagram of the rotating-tube phantom design with five smaller cylindrical samples along with temperature probe. (b) A photograph of the 5 tubes. (c) Magnitude images of the sagittal (one tube), axial, and coronal (multiple tube) views. (d) 3D rendering from MRI magnitude images.

### 2.1 Experimental Setup

The rotating-tube phantom consists of five cylindrical tubes (80mm length and 10mm outer diameter) alternated orthogonally along the central axis of a larger cylinder (610mm length and 140mm outer diameter) containing water (Fig. 1). Each tube contains one of the following: 0.5mM GdCl_3_, 1.0mM GdCl_3_, or 3.2mM CuSO_4_ (Fig. 1b). A rod extends from the internal rotation gears through the phantom and outside the MRI scanner, allowing the tubes to be manually rotated. Example MR images are shown for the three primary planes (Fig. 1c) and in a 3D rendering of the tubes (Fig. 1d). Temperature of the water was continuously monitored via a fiber optic probe (OpSens Medical, Québec, QC, Canada).

The paramagnetic salt solution Δ*χ* values were estimated from susceptibility theory and corrected using Curie’s law with the experimentally measured temperatures (21.0°C-22.0°C). Δ*χ* values were 0.336ppm and 0.168ppm for the 1mM GdCl_3_ and 0.5mM GdCl_3_, respectively, and 0.0804ppm for the 3.2mM CuSO_4_. Additional details on the calculation of Δ*χ* are in the Appendix.

### 2.2 Data Acquisition

MR data was collected on a 3T Siemens Biograph mMR (MR-PET scanner, Syngo MR E11 software) with a 6-element torso array and 9-element spine array coil for a total of 15 elements. To assess the reproducibility of phase estimation across pulse sequences, we acquired data with four Gradient echo (GRE) pulse sequences (details in Table 1):

- Single Echo GRE (SEGE): This is a commonly chosen protocol with QSM and other susceptibility-based techniques^28^, wherein a single echo time TE is measured as close to the *T*_2_* of tissue of interest (here, the target is 60ms for 1mM GdCl_3_). This maximizes the phase SNR at this *T*_2_*. In order to maximize the magnitude SNR at the chosen TE and TR, we set the readout bandwidth at its lowest possible value.
- Segmented Echo Planar Imaging (sEPI): A recently proposed sEPI sequence was shown to possess similar quality phase images as SEGE^29^, while acquiring full brain coverage much faster than SEGE. As with SEGE, phase images were generated at a single TE at the center of the echo train.
- Multi Echo GRE (MEGE): This protocol acquires multiple TEs in a single TR. The challenge with this technique is the choice of the echo spacing Δ*TE* and readout bandwidth BW. A short Δ*TE* reduces the likelihood of aliasing in the phase domain but introduces noise. A long Δ*TE* yields phase images with better SNR but suffers from potentially unrecoverable phase-aliasing errors. For example, in order to unwrap frequency offsets of ±150Hz, Δ*TE* must be less than 3.33ms. A common approach^28, 30^ is to acquire data with a short Δ*TE* and, in order to recover SNR efficiency similar to SEGE sequences^31^, acquire as many echoes as possible in a TR. However, due to hardware limitations, the readout bandwidth BW places a lower limit on Δ*TE*. In this work, we aimed to select the shortest Δ*TE* possible at the highest BW attainable with the MR system. This choice minimizes the likelihood of phase wrap errors which may not be recoverable by all phase unwrapping algorithms. We elaborate on this choice further in the Discussion section.
- MAGPI: This is an MEGE sequence that uses pre-optimized echo times and bandwidths selection that, when paired with a corresponding phase estimation algorithm, yields Maximum-Likelihood optimal phase estimates in the presence of wrapping, noise, and phase-offset errors^32, 33^.

**Table 1.** All data acquisition parameters.

| Sequence | Scan ID | TR (ms) | Echo Times (ms) | Bandwidth (Hz) | alpha (deg) | Resolution (mm<sup>3</sup>) | Acquisition Time (min:s) |
| --- | --- | --- | --- | --- | --- | --- | --- |
| **Single Echo GRE (SEGE)** | 1 | 25 | 16 | 80 | 15 | 1.0×1.0×2.0 | 1:59 |
|  | 2 | 35 | 25.7 | 80 | 15 | 0.5×0.5×2.0 | 4:35 |
|  | 3 | 45 | 30 | 80 | 15 | 1.0×1.0×2.0 | 3:35 |
| **Multi Echo GRE (MEGE)** | 1 | 25 | 2.5, 6.2, 9.9, 13.6, 17.3, 21.0 | 500 | 15 | 1.0×1.0×2.0 | 1:59 |
|  | 2 | 35 | 3, 7.8, 12.6, 17.4, 22.2, 27.0 | 530 | 15 | 0.5×0.5×2.0 | 4:35 |
|  | 3 | 45 | 2.5, 6.2, 9.9, 13.6, 17.3, 21.0, 24.7, 28.4, 32.1, 35.7, 39.5 | 500 | 15 | 1.0×1.0×2.0 | 3:35 |
| **sEPI** | 1 | 72 | 27, ETL = 15 | 430 | 20 | 1.0×1.0×2.0 | 0:31 |
|  | 2 | 72 | 27, ETL = 15 | 430 | 20 | 0.5×0.5×2.0 | 0:52 |
| **MAGPI** | 1 | 25 | 4.1, 8.9, 12.6, 16.3, 20.0 | 190 | 15 | 1.0×1.0×2.0 | 1:59 |
|  | 2 | 35 | 7.6, 13.0, 17.3, 21.5, 25.7, 30.0 | 250 | 15 | 0.5×0.5×2.0 | 4:35 |
|  | 3 | 45 | 6.6, 13.0, 18.5, 23.8, 29.0, 34.2, 39.4 | 200 | 15 | 1.0×1.0×2.0 | 3:35 |

All sequences were 3D excitations of a 128mm x 128mm x 128mm slab (64 slices). We used anisotropic voxels to boost SNR, a common practice for QSM and Susceptibility-Weighted Imaging^22, 23, 34^.

We assessed the effect of in-plane resolution and TR on phase estimate reproducibility with each protocol (Table 1). We also examined the reproducibility of phase estimates across nine different angles by advancing the apparatus approximately 18 degrees per turn.

### 2.3 Phase Estimation

Images generated in Steps 2-3 of QSM are commonly referred to as frequency, phase or field maps, depending on the units of the data. We interchangeably use these names in this work depending on context, and in our analysis, we convert all phase images to frequency via a simple scalar multiplication. Ten phase algorithms were selected to estimate the frequency offset image^32, 35-42^. Table 2 lists the methods used for each pulse sequence; multiple codes were downloaded from freely-available resources (e.g., MEDI) and integrated with the pipeline.

**Table 2.** All phase estimation methods. The first six methods were combined with other common phase processing techniques to process the multi-echo data, as described in the Methods section.

| Sequence | Method | Summary |
| --- | --- | --- |
| SEGE,<br>MEGE,<br>sEPI | **Unprocessed** | Default channel combined output phase image produced by the scanner |
|  | **Phun** | Region-based algorithm <sup>41</sup> |
|  | **GBC** | Goldstein's branch cut method <sup>37</sup> |
|  | **MEDI.RG** | Region growing algorithm from MEDI Toolbox <sup>40, 42</sup> |
|  | **MEDI.LP** | Laplacian algorithm from MEDI Toolbox <sup>39, 40</sup> |
|  | **Laplace** | Laplacian algorithm <sup>35</sup> |
| MEGE | **Slope** | Computes the frequency offset from the slope of the complex data across echoes <sup>36</sup> |
|  | **Division** | Computes the frequency offset from the complex division of the data at successive echoes <sup>38</sup> |
|  | **MAGPI-unopt** | Applies the MAGPI post-processing algorithm using a random (un-optimized) subset of echoes from the MEGE sequence. Since MAGPI requires uneven echo spacing, only four unequally spaced echoes were selected. <sup>32</sup> |
| MAGPI | **MAGPI** | Maximum likelihood estimate of phase from optimal echo spacing <sup>32</sup> |

All phase estimation methods were applied with default parameters in 3D over the entire acquisition volume. Apart from the MAGPI algorithm, which operates on raw k-space channel-uncombined data, all methods (including MAGPI-unopt) operated on unprocessed phase data obtained using the vendor-provided adaptive-coil-combine method. Each unwrapping method used the SNR in the magnitude image to guide unwrapping orientation in the phase domain: e.g., the Laplacian-based methods used this SNR to mask the entire image, while others (region-growing, GBC) masked the phase values in regions with poor SNR.

For MEGE sequences, the multi-echo data is processed using the following five categories of algorithms:

1. Spatial phase unwrapping at each echo, followed by temporal combination of the resulting images using a weighted averaging method that is phase SNR-optimal^43^

2. Spatial phase unwrapping at each echo with weighted averaging (as in 1), but a 1D phase unwrapping step is used just before weighted averaging. This is meant to correct any remaining aliasing that spatial unwrapping failed to correct.

3. Direct temporal phase estimation (Slope, Division) applied in complex domain. These methods correctly unwrap the phase over time, provided the inherent frequency is less than the Nyquist frequency associated with the echo spacing.

4. Temporal phase combination (as in 3), followed by 3D spatial phase unwrapping to correct errors encountered with temporal phase estimation.

5. Maximum Likelihood-based combination of multi-echo and multi-channel data (MAGPI)^32^. This method solves the phase estimation problem on a voxel-by-voxel basis, without resorting to spatial averaging techniques.

All phase images are eventually converted to frequency offset (Hz) by dividing by (2π x phase evolution time).

### 2.4 Adjusting for field due to rotation of apparatus

In order to accurately estimate Δ*χ* from the phase images, we need to remove the global field effects resulting from the tubes rotating in the magnetic field. We call this process “frequency referencing”.

The complete field inside a voxel can be written as:

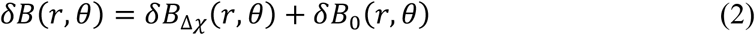

where *r* is the spatial coordinate of the voxel and *θ* is the rotation angle of the tube relative to *B, δB*_Δ*χ*_(*r,θ*) is the field caused by magnetic susceptibility variations (such as the one due to a homogeneous cylindrical object immersed in a homogeneous sphere), and *δB*_0_(*r,θ*) is an unknown frequency offset component. We separate *δB*_0_(*r,θ*) into a component that varies only spatially and a component that varies only with the rotation of the apparatus:

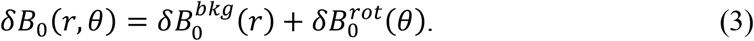

At a given angle of rotation, sources of spatially varying global offsets 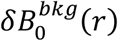 are field inhomogeneity (imperfections of magnetic field/coils), bulk magnetic susceptibility of the apparatus^44, 45^ and coil phase offset^46^. As the apparatus is rotated, in the absence of “shimming” at the console, the center frequency will be shifted due to the bulk magnetic susceptibility of the entire apparatus^47, 48^. Our goal is to extract *Δ*χ by fitting *δB*(*r,θ*) to the angle of rotation *θ*, inside the tube. 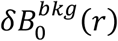 is a nuisance parameter that can be easily accounted for during the fitting process by allowing for a constant shift to the cosine.

First, we compute an estimate of 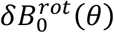 using the average frequency in a static region outside the “Tube+Sphere” system (Fig. 2a). The average (indicated by < >) is taken over pixels in a region *r*_*out*_ distant from local susceptibility effects:

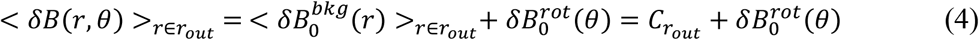

where we define 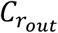 to be a variable that is only a function of the referencing region. Then, the referencing step consists of:

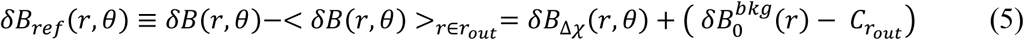

thus removing 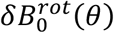. Our goal is to use the referenced field, *δB*_*ref*_(*r,θ*), at every pixel in the tube center, *r* = *r*_*in*_, to fit the field variation to the angle of rotation *θ*. The only component that varies with *θ* is the first term on right side of Eq 5. The estimation step is a simple fit with respect to *θ*, along with an arbitrary shift for the constant: 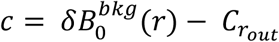. For the case of a cylinder (Eq 1), the estimate can be obtained by solving:

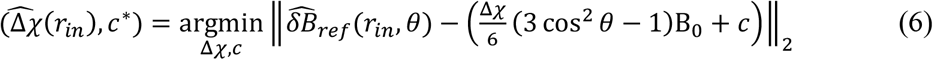

Fitting c effectively amounts to shifting the midline of the data to match the model (across all angles). We used a bisquare-weighting method to fit this midline. We include a few examples of 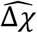 estimation using Eq 6 in Table A.1.

**Fig. 2.**
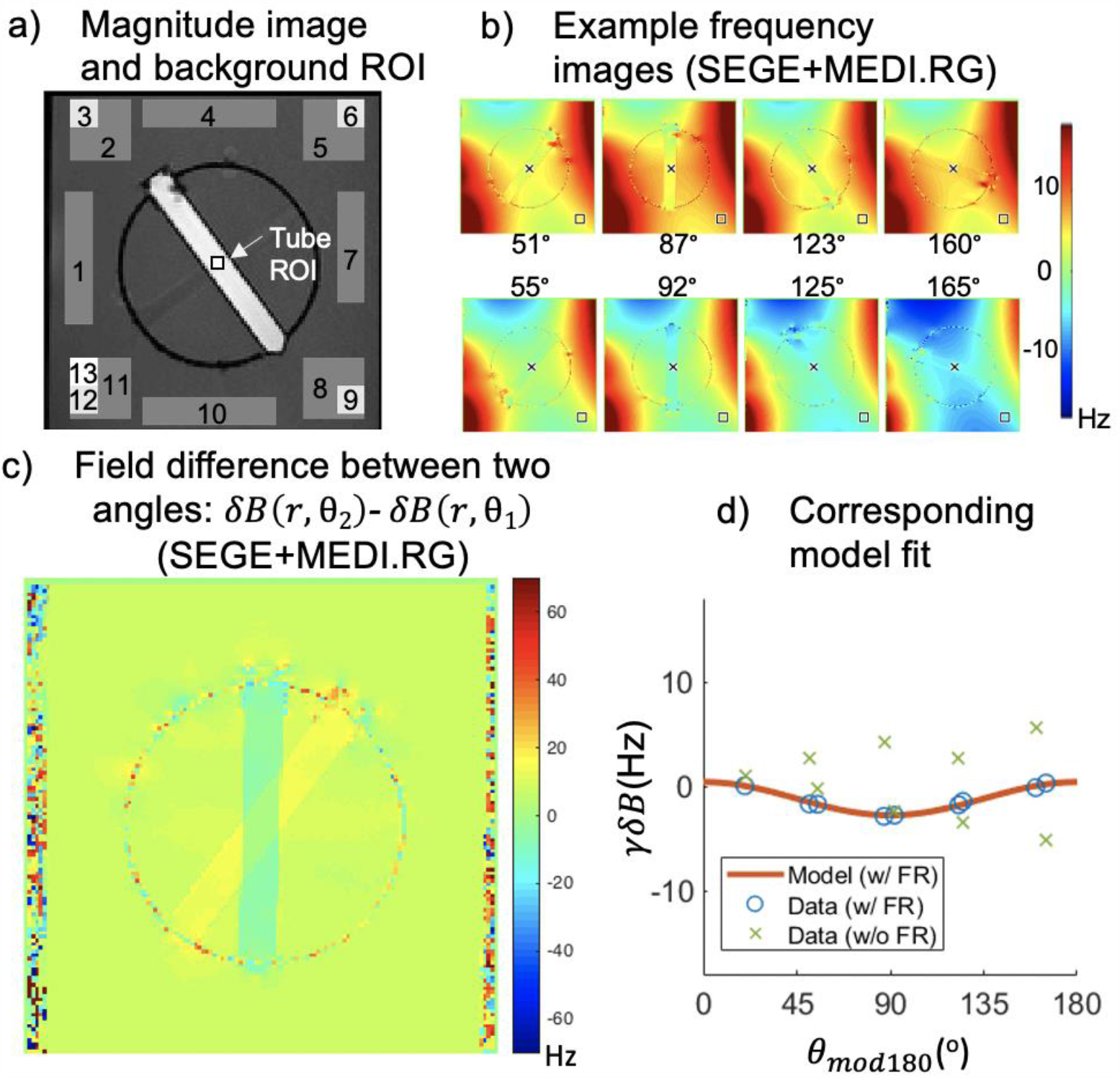
(a) An example magnitude GRE image showing both the tube ROI (small black box in center) used for the analysis as well as the placement of the 13 background ROIs. (b) Example frequency images in tube 3 (0.5 mM GdCl3) obtained with the Single GRE sequence (1 mm, TR 45 ms) and reconstructed with MEDI.RG phase estimation method. (c) Also using SEGE+MEDI.RG, the field difference between two angles (***δB*_*total*_(*r*,θ2)-*δB*_*total*_(*r*,θ1)**) showing that this difference is attributable to the spatially invariant component (in homogeneous areas) and a spatially variant component (in areas close to material boundaries). The spatially invariant component of the field difference is removed with frequency referencing. (d) A plot of the frequency against the angle of rotation (modulo 180°) is shown for each of the data without frequency referencing (in green x), data after frequency referencing using the frequency reference ROI #9 (blue circles), and the model’s prediction of the frequency (solid red line).

Figure 2a shows a magnitude GRE image of tube 3 (0.5mM GdCl_3_) in one orientation. Figure 2b shows frequency offset images corresponding to different rotations. Figure 2c shows a typical field difference between two angles (*δB*(*r*,θ2)*-δB*(*r*,θ1)). According to Eq 3, this difference is equal to: 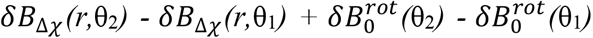, predicting spatial variations only in locations close to areas with susceptibility changes and a constant field in homogeneous locations. This is precisely what we observe in Fig. 2c. Figure 2d shows a plot of the frequency values for a voxel inside the tube prior to frequency referencing (symbol ‘x’). The resulting plot does not follow the expected sinusoid (solid line). After applying our proposed frequency referencing step, we observe the expected sinusoidal shape (symbol ‘o’). We apply this frequency referencing method after each phase estimation algorithm. To investigate repeatability with respect to the location of frequency reference estimate, we apply this process in 13 different regions selected across static areas of the phantom (Fig. 2a).

### 2.5 Error Analysis

We use the theoretically-determined values of Δ*χ* to predict the field values at each angle that would have been measured with ideal methods in Steps 1 & 2. We then compute the error (Hz) between measured frequency 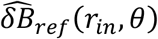 and expected frequency offset:

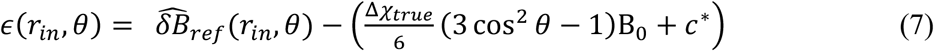

where *c*^*\**^is obtained by solving Eq 6 with Δ*χ* set to Δ*χ*_*true*_.

To account for any dependence on tube content, we compute the absolute relative error:

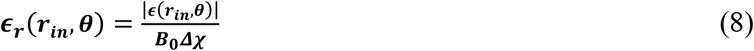

where the absolute value is used instead of the signed error due to the irrelevance of sign in this context.

Error statistics were computed for each voxel in the tube ROI (mean tube ROI sizes were 17 pixels at 1mm resolution and 57 pixels at 0.5mm resolution), for each slice (2 slices per tube), each tube (5 total), each angle (9 total rotations), using each applicable phase estimation method (from a possible 10), each background phase removal ROI (13 total), and each sequence with its respective resolution and TR variations (11 total). Cumulatively, 2.36 million frequency values were analyzed in this experiment.

The large number of data points allows us to extract statistics about *ϵ* and *ϵ*_*r*_ including their probability mass function, Pr(*ϵ*) and Pr(*ϵ*_*r*_),shown in Fig. 3a and 3b for an example method. The probability of *ϵ* for this example exhibits a multi-modal distribution, and therefore is not a Gaussian. While we can report the absolute bias and standard deviations from such a distribution (Fig. 3), it would not be descriptive of Pr(*ϵ*). A more practical measure is the likelihood of observing absolute relative errors less than or equal to a threshold, *τ*. This is obtained by integrating Pr(*ϵ*_*r*_) between 0 and *τ*,a measure known as the Cumulative Distribution Function (CDF),

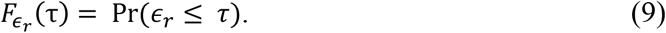

For example, we can see from the CDF in Fig. 3c that the probability of observing errors less than 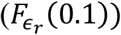 is about 66%. The CDF can be used to capture phase errors that are dominated by outliers, as well as phase errors that result from generally poor/unreliable model fitting. The ideal CDF is a step function, and any presence of outliers/large errors yields a CDF with slow convergence to 1. The frequency of large errors is seen from the magnitude of the deviation of the CDF from 1.0 at any given threshold.

**Fig. 3.**
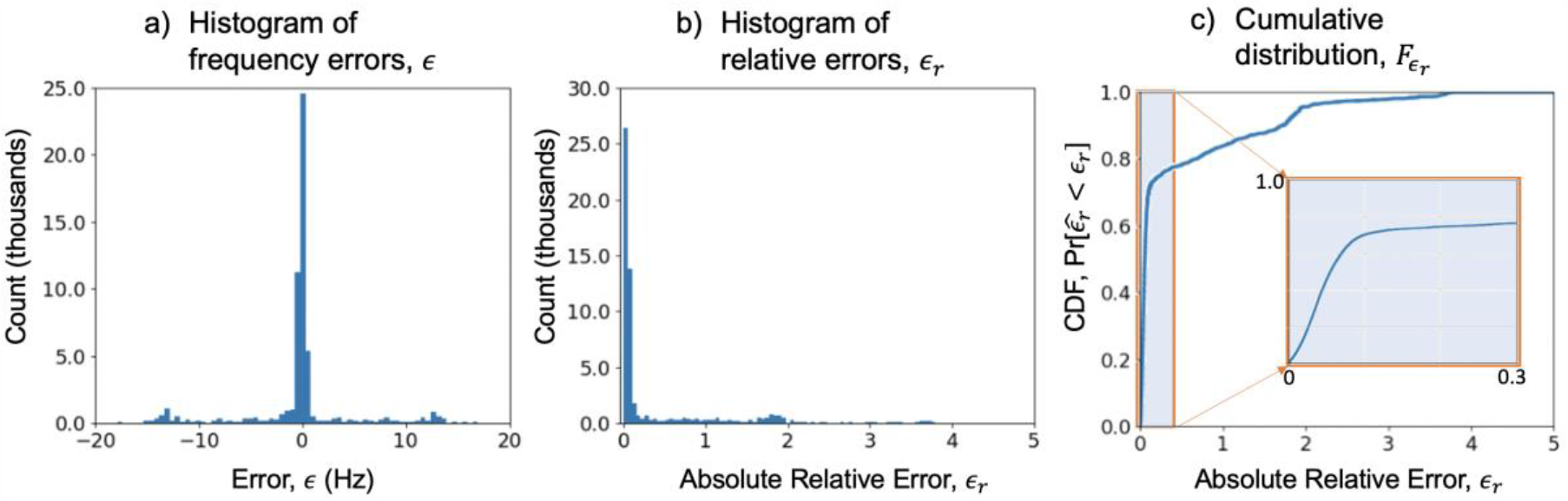
(a) The histogram of the error ***ϵ*** (in Hz) when the measurement is obtained with the SEGE+GBC phase estimation method. The error data is pooled over all ROI voxels, all backgrounds, rotations and tubes. (b) The histogram of the absolute relative error ***ϵ*_*r*_** seen with the same measurement. (c) The corresponding Cumulative Distribution Function 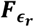 of the error in (b). The inset figure shows a portion of this CDF within [0,0.3] to better illustrate the distributions within reasonable error range. In this case, the CDF shows that the probability of obtaining ***ϵ*_*r*_** less than 0.1 is around 0.66 with the SEGE+GBC pair. The maximum observed ***ϵ*** was 27.0 Hz, and the maximum observed ***ϵ*_*r*_** was 6.85, illustrating the occasional large errors that may result with SEGE+GBC.

## 3 Results

Table 3 summarizes the error statistics for all combination of sequences and algorithms. The first column lists the sequence type, and the second and third column indicate the name and category of each post-processing algorithm, respectively. We report the mean and standard deviations of both *ϵ* and *ϵ*_*r*_. We also report 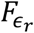 (0.1) pooled over all background ROIs, as well as the worst and best case 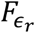(0.1) encountered in those ROIs. These quantities are computed from data that includes all tubes and angles. A representative subset of these results is selected for more detailed analysis and illustration in Fig s4-6.

**Table 3.** Error statistics for each of the estimation methods and acquisition protocols. The mean and standard deviation of error measurements *ϵ* (Hz), as well as the absolute relative errors *ϵ*_*r*_, are provided in columns 4-7. Columns 8-9 show 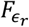 (0.1) of the overall data, as well as the range (min and max) of 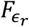 (0.1) across reference backgrounds.

| Prot-ocol | Post-Processing | Method | $\mu_{\epsilon}$ (Hz) | $\sigma_{\epsilon}$ (Hz) | $\mu_{\epsilon_r}$ | $\sigma_{\epsilon_r}$ | $F_{\epsilon_r}(0.1)$ | $[F_{\epsilon_{rw}}(0.1), F_{\epsilon_{rb}}(0.1)]$ |
| --- | --- | --- | --- | --- | --- | --- | --- | --- |
| SEGE | Spatial 3D unwrap | Unpro-cessed | 0.29 | 3.85 | 0.52 | 0.92 | 0.48 | [0.24, 0.76] |
|  |  | Phun | 0.21 | 5.12 | 0.35 | 1.38 | 0.67 | [0.49, 0.89] |
|  |  | GBC | -0.22 | 4.83 | 0.53 | 1.08 | 0.66 | [0.47, 0.82] |
|  |  | MEDI.RG | -0.19 | 1.60 | 0.08 | 0.22 | 0.90 | [0.84, 0.96] |
|  |  | MEDI.LP | -0.01 | 1.07 | 0.14 | 0.12 | 0.50 | [0.38, 0.62] |
|  |  | Laplace | 0.00 | 1.01 | 0.14 | 0.15 | 0.54 | [0.40, 0.66] |
| sEPI | Spatial 3D unwrap | Unpro-cessed | -0.34 | 4.19 | 0.58 | 0.76 | 0.43 | [0.32, 0.52] |
|  |  | Phun | -0.10 | 5.23 | 0.73 | 1.30 | 0.45 | [0.31, 0.56] |
|  |  | GBC | -0.32 | 3.80 | 0.42 | 0.97 | 0.66 | [0.31, 0.91] |
|  |  | MEDI.RG | 1.03 | 4.14 | 0.32 | 1.15 | 0.85 | [0.79, 0.91] |
|  |  | MEDI.LP | 0.00 | 1.14 | 0.12 | 0.12 | 0.57 | [0.51, 0.65] |
|  |  | Laplace | 0.00 | 0.95 | 0.12 | 0.11 | 0.58 | [0.47, 0.68] |
| Multi Echoes | 1. Spatial 3D unwrap → Temporal average | Unpro-cessed | 0.01 | 2.60 | 0.48 | 0.41 | 0.16 | [0.08, 0.23] |
|  |  | Phun | 0.39 | 4.75 | 0.30 | 1.23 | 0.57 | [0.48, 0.69] |
|  |  | GBC | 0.31 | 2.21 | 0.15 | 0.36 | 0.78 | [0.59, 0.91] |
|  |  | MEDI.RG | -0.37 | 3.47 | 0.66 | 0.95 | 0.35 | [0.09, 0.63] |
|  |  | MEDI.LP | -0.01 | 0.89 | 0.11 | 0.09 | 0.58 | [0.49, 0.67] |
|  |  | Laplace | 0.00 | 0.82 | 0.12 | 0.12 | 0.58 | [0.44, 0.68] |
| | 2. Spatial 3D unwrap → Temporal 1D unwrap → Temporal average | Phun | 0.19 | 7.66 | 0.94 | 2.23 | 0.55 | [0.12, 0.96] |
|  |  | GBC | 0.20 | 7.78 | 0.96 | 2.25 | 0.55 | [0.12, 0.96] |
|  |  | MEDI.RG | 0.13 | 7.95 | 0.97 | 2.29 | 0.55 | [0.12, 0.96] |
|  |  | MEDI.LP | -0.01 | 0.89 | 0.11 | 0.09 | 0.58 | [0.49, 0.67] |
|  |  | Laplace | 0.00 | 0.82 | 0.12 | 0.13 | 0.58 | [0.45, 0.68] |
|  | 3. Temporal estimation | Slope | 0.26 | 9.28 | 1.15 | 2.79 | 0.55 | [0.13, 0.96] |
|  |  | Division | 0.26 | 9.27 | 1.15 | 2.78 | 0.55 | [0.13, 0.95] |
| | 4. Temporal estimation → Spatial 3D unwrap | Phun | 0.04 | 11.04 | 1.17 | 3.02 | 0.53 | [0.06, 0.95] |
|  |  | GBC | 0.17 | 9.18 | 1.10 | 2.69 | 0.54 | [0.09, 0.95] |
|  |  | MEDI.RG | 0.47 | 62.07 | 10.11 | 16.54 | 0.21 | [0.00, 0.76] |
|  |  | MEDI.LP | 0.16 | 3.87 | 0.54 | 1.16 | 0.43 | [0.20, 0.70] |
|  |  | Laplace | 0.22 | 6.89 | 0.89 | 2.05 | 0.36 | [0.10, 0.66] |
|  | 5. MAGPI | MAGPI unopt | 0.00 | 0.30 | 0.05 | 0.05 | 0.90 | [0.78, 0.97] |
|  |  | MAGPI | 0.00 | 0.27 | 0.05 | 0.04 | 0.91 | [0.84, 0.98] |

**Fig. 4.**
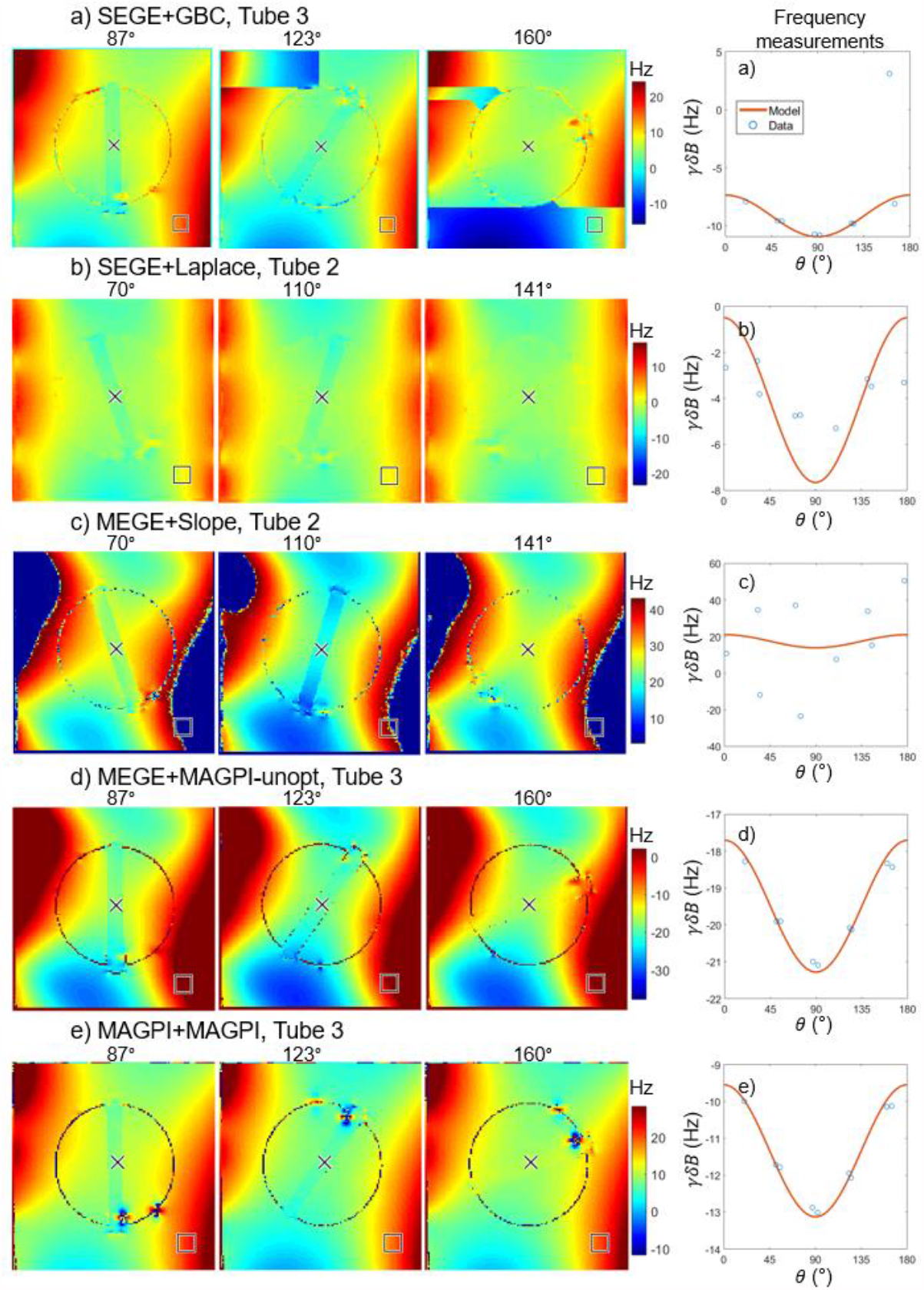
Each row in this figure shows five different examples of sequence (at 1 mm resolution, TR = 45 ms) and phase estimation method pairs for three different angles of rotation in different tubes. These examples were selected to illustrate the spatial nature of different phase estimations errors and artifacts. The corresponding plots in the last column show the resulting frequency measurements as a function of angle of rotation (modulo 180°), for a voxel inside the tube, after frequency referencing (blue circle). The predicted frequency offset as obtained from Eq 1 is also shown in solid red line. The background ROI (#9) used is shown with a square overlaid on the frequency maps. (a) SEGE+GBC shows a phase wrapping error in tube 3 in the frequency reference ROI for the 160 degree angle. (b) SEGE+Laplace shows smoothly varying frequency maps; however, the values deviate from the expected result at every angle. (c) MEGE+Slope, without 3D phase unwrapping, shows the presence of phase wrapping in areas with large frequency values. The frequency reference ROI is within a phase wrapped region across all angles. (d-e) MEGE+MAGPI-unopt and MAGPI’s phase estimation show frequency maps consistent with values predicted from the model.

**Fig. 5.**
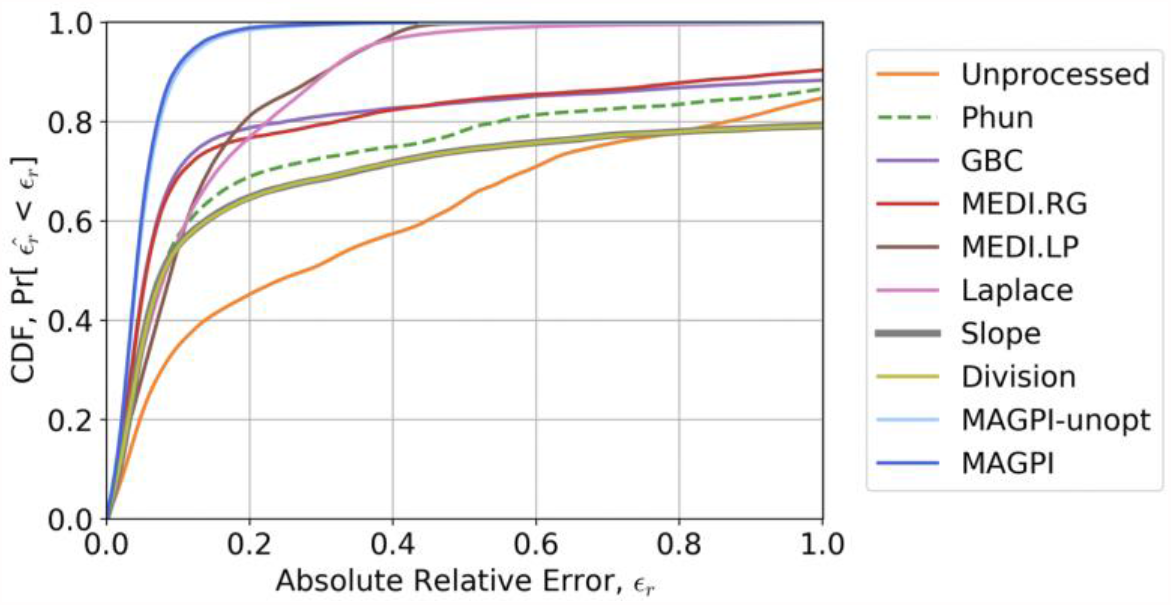
CDF *F*_*ϵ*_*r*of the absolute relative error obtained for each of the phase estimation methods studied in this work, when the error is pooled over all sequences, slices, tubes, angles and frequency referencing ROIs. Note that some methods do not converge to probability of 1, due to the presence of errors greater than 100%.

**Fig. 6.**
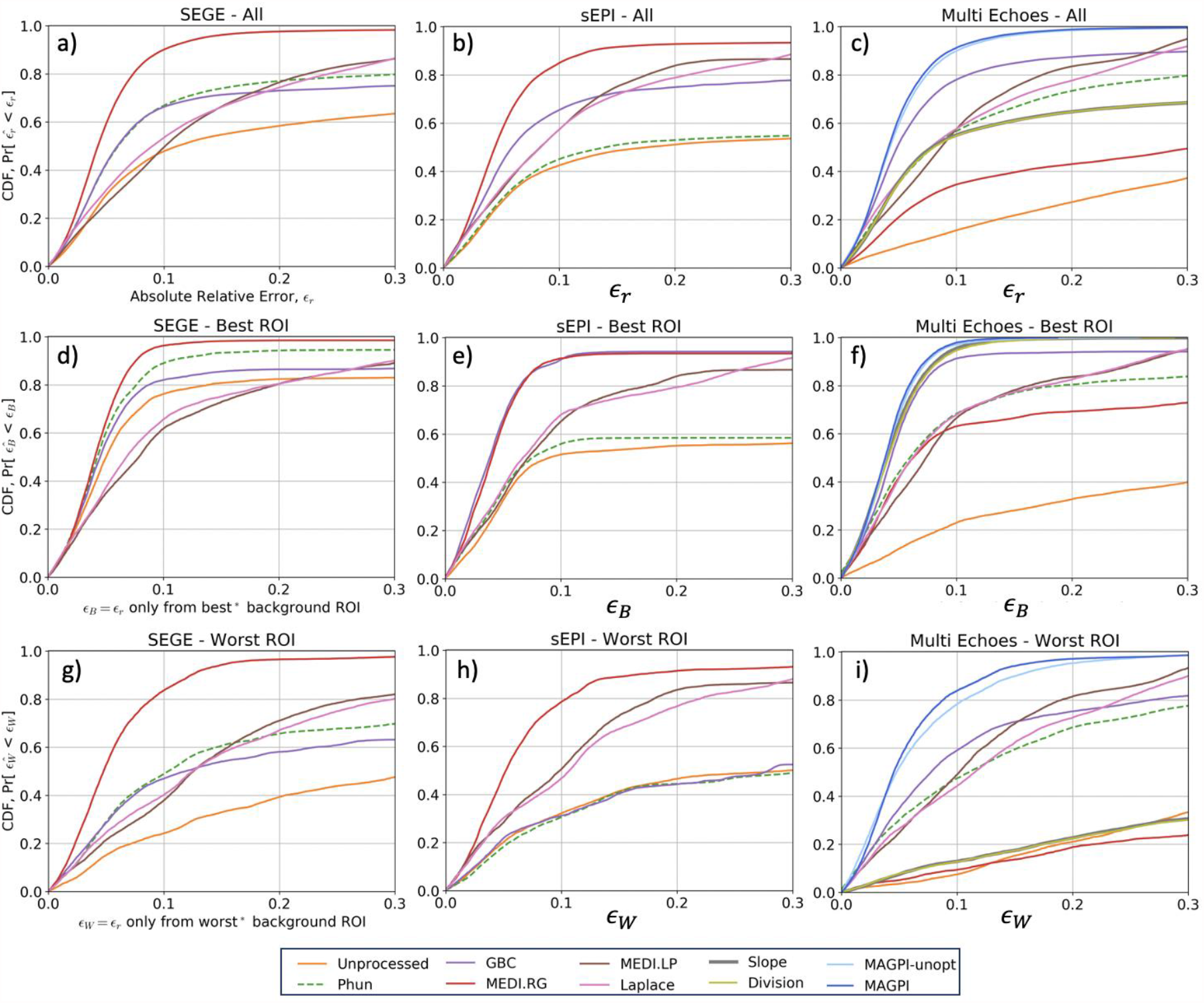
The CDFs for each acquisition and phase estimation method pair. Each column corresponds to a different type of acquisition: single echo acquisitions (SEGE) are shown in the first column, accelerated acquisitions with sEPI in the second, and multi echo acquisitions (MEGE & MAGPI) in the third. The first row (a-c) shows the CDF when all the error data is pooled, over all voxels, slices, tubes, and background ROIs. Given the large variability in performance for different background ROIs, we show in the second (d-f) and third (g-i) rows the CDFs obtained in the “best” and “worst” ROIs, respectively. A highly reproducible method will have a similar curve shape across all plots.

Figure 4 shows a subset of frequency-offset images for different sequence+algorithm pairs, at 3 of the 9 angles of rotation. This figure illustrates typical challenges with phase estimation methods. For example, in Fig. 4a, we observe phase unwrapping errors in SEGE+GBC, with abrupt jumps across contiguous regions. The corresponding frequency vs angle plot (last column in Fig. 4a) shows that incorrect frequency referencing in these areas (square in figure) yield occasional mismatch between measurement and predictions at certain angles. SEGE+Laplace demonstrates a smoothly varying frequency map across the FOV; however, the resulting data deviates from the expected theoretical values at almost every angle (Fig. 4b). MEGE+Slope, a direct temporal phase estimation method (MEGE category 3), exhibits phase wrapping errors when the underlying frequency value is larger than the bandwidth allowable by the Δ*TE* (Fig. 4c). Placing a frequency referencing ROI in these areas yields incorrect values at the respective angles. Note that the example frequency reference ROI (#9 in Fig. 2) is meant to highlight the phase errors or artifacts observed. Figures 4d and 4e show that the results from MAGPI-unopt and MAGPI are consistent with those predicted from theory.

The CDF of *ϵ*_*r*_ collects the errors, such as those observed in Fig. 4, over a variety of acquisition and processing parameters. Figure 5 shows 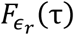 for all algorithms when *ϵ*_*r*_ is pooled over all voxels, background ROIs, slices, tubes, and sequence variations. This represents an overall summary of algorithm behavior, irrespective of which parameter was used in acquisition and post-processing. We see that MAGPI attains a nearly ideal CDF, with 0.91 probability of relative errors, *ϵ*_*r*_, less than 0.1 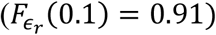 and rapidly converges to 1 (Fig. 5). MAGPI and MAGPI-unopt achieve similar CDFs, with MAGPI performing slightly better, as expected. MEDI-RG and GBC phase unwrapping methods, both based on region growing, have similar CDFs, with 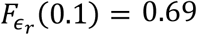 and 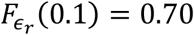, respectively. The unprocessed phase images have the most artifacts and, thus, the lowest CDF across all *ϵ*_*r*_. Figure 5 focuses on the CDF for *ϵ*_*r*_ in [0,1.0] to highlight the different convergence pattern (distribution/frequency of errors) in that domain. The CDF extends beyond *ϵ*_*r*_ = 1.0 for any occurrence of relative errors greater than 100%.

Next, we explore the behavior of 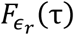 as a function of data acquisition strategy. In Figs. 6a-c, we group results by three sequence types: SEGE, sEPI and MEGE. Since MAGPI is a multi-echo sequence, we include MAGPI in the MEGE category. For each CDF curve 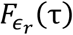, *ϵ*_*r*_ is pooled across all pixels, background ROIs, slices, tubes, and variations of TR and resolution within that sequence type. We also explore variability of 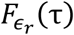 with the frequency referencing method. For each sequence, we show CDFs in the “best ROI” (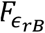,Figs 6d-f) and “worst ROI” (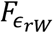, Figs 6g-i). Here, “best/worst” ROI is defined as the one having the largest/smallest 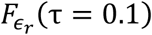 for a given dataset. The separation between 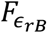 and 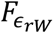 demonstrates the robustness (or lack thereof) of a method to frequency reference ROI selection.

## 4 Discussion

A reliable pulse sequence protocol, with repeatable and reproducible phase estimation, is necessary step to develop robust QSM methods for clinical use. Previous work examined the reproducibility of certain QSM methods using phantoms^18, 21, 49^, simulation^18, 21, 50^, and human subjects^15, 50^. We used a rotating-tube phantom to quantitatively evaluate methods used for data acquisition and phase estimation. This is not the first study using long tubes nor is it the first to position tubes relative to B_0_; however, compared to previous work^18, 20, 21, 49, 51-53^, more acquisition and phase estimation methods were considered. Specifically, we used ∼90 different combinations of pulse sequences and phase estimation methods to analyze millions of measurements from different ROIs, tube contents, rotations and sequence parameters.

Our analysis considered diverse scenarios encountered in QSM applications. We sampled errors in different frequency reference ROIs, because accurate field estimates everywhere in the FOV are important for Steps 3 and 4 of the QSM process. We sampled errors across different tube contents to capture any variability with chemical content. We attempted to capture the more common sequences in QSM, though we could not include all. MAGPI is not a common sequence/method (and is not publicly available) but was used due to its potential to overcome some phase estimation challenges. We introduced scan variability using different TRs and resolutions. While other variations on scan parameters exist, it is not feasible to collect every variation. We attempted to use each phase estimation method available to us and included unprocessed phase data to demonstrate baseline performance of validation criteria. Additional phase estimation methods could be retrospectively used on the data set, which we aim to make publicly available. Finally, we rotated the tubes and apparatus to capture different geometric orientations with respect to B_0_. We used rotation angle, in addition to knowledge of the sample’s magnetic susceptibility, with the closed-form solution given by Eq. 1 to derive predictions of the expected phase at every angle. This vast amount of data ultimately allowed us to estimate the probability distribution of field/phase error with every QSM method, along with other important statistics.

Table 3 displays error metrics for all combinations of sequences (Step 1) and methods (Step 2). Our results showed varying degrees of accuracy and precision over all tested methods. For example, while the majority of methods resulted in *μ*_*ϵ*_ less than 1Hz (note the particularly small *μ*_*ϵ*_ with MEDI-LP, Laplace, MEGE+MAGPI-unopt and MAGPI), the only methods with 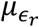 are SEGE+MEDI.RG (7.6%), MEGE+MAGPI-unopt (5.3%), and MAGPI (5.0%). We observed a similar trend with precision, whereby methods with the lowest relative standard deviation 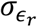 were MEGE+MEDI.LP (8.9%), MEGE+MAGPI-unopt (4.8%), and MAGPI (4.1%). The detailed behavior of the error is captured by the CDF (or PDF) of the data (Fig. 5). A summary of the CDF is in the second-to-last column of Table 3 where we show the probability of observing relative errors <10%, 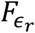 (10%), which captures the frequency *relatively* acceptable errors occur. Almost always, 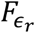 (10%) of Laplace-based methods was amongst the worst performers, resulting in acceptable errors only 50-60% of the time. Laplace-based methods had smooth phase maps with qualitatively no apparent phase jumps. However, analysis showed that Laplace phase images result in quantitatively larger errors than other methods, suggesting incorrect phase unwrapping results, similar to Chen et al^54^. Other methods with poor (low) 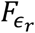(10%) were the unprocessed phase data, MEGE+Slope/Div and MEGE+MEDI.RG, which had large phase-unwrapping errors in a significant proportion of the data. 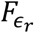(10%) is an arbitrary point at which we highlight the behavior of 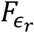 and does not represent the entirety of the distribution of error (or CDF). For example, 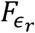(10%) of SEGE+MEDI.RG was comparable to MEGE+MAGPI-unopt and MAGPI, despite the comparatively poorer (larger) 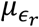 and 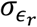 of SEGE+MEDI.RG. This is due to relative errors falling mostly within the chosen 10% threshold for these methods. Nevertheless, we report 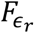(10%) instead of plotting CDF for every sequence+method pair.

We explored the dependence of errors on frequency reference ROI location (Fig. 6). Since phase estimation errors (particularly large errors) are undesirable anywhere in the FOV, any spatial variation of the CDF highlights the potential dependence of the method on user-intervention and/or its automated processing. We show the range of 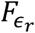(10%) observed across the 13 different frequency reference ROIs in the last column of Table 3. The results suggest that the most repeatable methods across background ROIs are SEGE+MEDI.RG, sEPI+MEDI.RG, MEGE+MAGPI-unopt, and MAGPI.

The MEGE data in particular was processed using four broad classes of post-processing algorithms. We note the following about these algorithms:

i. The results obtained with methods in Categories 1 & 2 were fundamentally similar.That is, additional 1D-temporal processing does not alter the performance of 3D spatial unwrapping methods (with the exception of Laplacian-based methods, which we discuss below). This redundancy is due to the inherent Nyquist limitation associated with the echo spacing. As a result, we focus on the distinctive results of Category 1: spatial phase unwrapping + weighted averaging of echoes (shaded MEGE rows of Table 3).
ii. Some post-processing methods used in MEGE Categories 1 & 2 performed more poorly with MEGE than with SEGE (e.g., MEDI-RG). We believe this is due to the relatively larger bandwidth used with MEGE acquisitions, which result in noisier images at each echo. Higher BW acquisitions were needed with MEGE to accommodate temporal methods (MEGE Category 3 & 4), which require short echo spacing. It is possible that MEGE+MEDI-RG would perform better with lower BW (wider echo spacing). Due to time/complexity constraints, we were unable to explore every possible MEGE variation that favors specific algorithms. This is a limitation of this study.
iii. MEGE Category 3 methods (Division/Slope) were straightforward to apply but resulted in a wide range of errors. This is due to large errors observed in frequency reference ROIs where the underlying frequency-offset value is larger than what is allowable by the smallest echo spacing. While such errors are avoidable with shorter echo spacing, this is not always possible (as was the case here) due to hardware constraints on readout bandwidth, resolution, FOV, etc. While Slope and Division are straightforward to apply, they result a sub-optimal combination of echoes, with noisy phase estimates.
iv. MEGE Category 4 methods have a similar performance to Category 3 methods. That is, spatial phase unwrapping did not seem to markedly improve the performance of temporal phase unwrapping. This is potentially due to the hard-to-unwrap noisy boundary lines observed with Category 3 methods, as shown in MEGE+Slope example in Figure 4, that are still present with Category 4 methods.

Finally, we explored the dependence of the CDF on scan variability and tube contents (Table 4). 4). Because the performance of some methods is dominated by frequency reference ROI (seen in Table 3, Fig. 6), Table 4 shows results for the “best ROI” for a given sequence+method pair. Here, “best ROI” is the largest 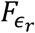(10%) observed over all frequency reference ROIs, for every method, sequence, scan variation, tube and slice. We note the following from this table:

1. Not all sequence+method pairs were invariant to scan and/or tube content. sEPI+Phun, and sEPI+Unprocessed had greatest variability, followed by all of the Laplace-based methods (LP, MEDI.LP); indicating that some methods produced inconsistent phase even in their best-case scenario. Laplace-based techniques generated smoothly-varying, though numerically inaccurate phase maps, irrespective of sequence, and sEPI+Phun suffered from sequence-dependent errors.
2. Considering the best-case ROI scenario, we did not observe consistent performance differences between pairing methods with either SEGE or sEPI. The sEPI sequence provides a significant acceleration over SEGE via its segmented GRE approach and performed better with some methods (GBC) and worse with others (Phun).
3. Table 4 compares Sequence+Method pair performance as a function of variability of scan parameters. For example, MEGE+Slope performed consistently well 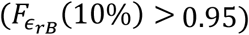,irrespective of the scan type used; while SEGE+Laplace struggled to estimate the correct phase, irrespective of scan. SEGE+GBC, though, only struggled with Scan 2, which had a shorter TR than Scan 1.
4. The same analysis can be applied to tube contents. MAGPI achieved 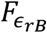(%) close to 1.0 for all tubes and 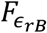(%) = 0.91 in Tube 5. sEPI+Phun had inconsistent performance across tubes of similar content (Tubes 1&2 and 3&4). Among the best performing methods, Tube 5 exhibited lower 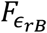(%) compared to other tubes, which could be due to challenges defining Δ*χ*_*th*_ and the complexity of CuSO_4_ compared to GdCl_3_.

**Table 4.** Probability of relative errors < 0.1 in the “best” background ROI, i.e., 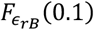,across sequences, scan variability (ID is as listed in Table 1) and phantom tube components. The tube contents are 1.0 mM GdCl_3_ in tubes 1 and 2, 0.5 mM GdCl_3_ in tubes 3 and 4, and 3.2 mM CuSO_4_ in tube 5. The last column is the standard deviation of all 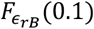 for the given method.

| Method | Sequence | Scan ID 1 | Scan ID 2 | Scan ID 3 | Tube 1 | Tube 2 | Tube 3 | Tube 4 | Tube 5 | std $F_{\epsilon_{rB}}(0.1)$ |
| --- | --- | --- | --- | --- | --- | --- | --- | --- | --- | --- |
| Unprocessed | SEGE | 0.79 | 0.86 | 0.47 | 0.59 | 0.88 | 0.83 | 0.89 | 0.68 | 0.32 |
|  | MEGE | 0.25 | 0.21 | 0.27 | 0.25 | 0.23 | 0.25 | 0.26 | 0.18 |  |
|  | sEPI | 0.51 | 0.55 | - | 1.00 | 0.00 | 0.86 | 0.06 | 0.77 |  |
| Phun | SEGE | 0.91 | 0.96 | 0.68 | 0.98 | 0.88 | 0.83 | 0.87 | 0.88 | 0.24 |
|  | MEGE | 0.67 | 0.73 | 0.64 | 0.89 | 0.51 | 0.54 | 0.66 | 0.80 |  |
|  | sEPI | 0.56 | 0.58 | - | 0.92 | 0.00 | 0.87 | 0.37 | 0.79 |  |
| GBC | SEGE | 0.96 | 0.73 | 0.96 | 0.75 | 0.79 | 0.97 | 0.93 | 0.73 | 0.08 |
|  | MEGE | 0.95 | 0.88 | 0.96 | 0.89 | 0.93 | 0.97 | 1.00 | 0.80 |  |
|  | sEPI | 0.89 | 0.97 | - | 0.95 | 0.89 | 0.90 | 1.00 | 0.82 |  |
| MEDI.RG | SEGE | 0.97 | 0.96 | 0.97 | 1.00 | 0.94 | 1.00 | 1.00 | 0.89 | 0.20 |
|  | MEGE | 0.24 | 0.76 | 0.65 | 0.63 | 0.61 | 0.64 | 0.67 | 0.63 |  |
|  | sEPI | 0.91 | 0.92 | - | 1.00 | 0.84 | 0.94 | 0.97 | 0.84 |  |
| MEDI.LP | SEGE | 0.63 | 0.63 | 0.59 | 0.36 | 0.58 | 0.53 | 0.96 | 0.70 | 0.18 |
|  | MEGE | 0.68 | 0.67 | 0.64 | 0.36 | 0.75 | 0.53 | 0.95 | 0.76 |  |
|  | sEPI | 0.65 | 0.66 | - | 0.36 | 0.63 | 0.53 | 1.00 | 0.78 |  |
| Laplace | SEGE | 0.57 | 0.68 | 0.68 | 0.36 | 0.97 | 0.52 | 0.85 | 0.56 | 0.19 |
|  | MEGE | 0.62 | 0.72 | 0.64 | 0.36 | 1.00 | 0.69 | 0.90 | 0.46 |  |
|  | sEPI | 0.66 | 0.74 | - | 0.40 | 0.93 | 0.58 | 0.89 | 0.60 |  |
| Slope | MEGE | 0.96 | 0.95 | 0.97 | 0.99 | 1.00 | 0.95 | 0.99 | 0.82 | 0.06 |
| Division | MEGE | 0.95 | 0.94 | 0.96 | 0.99 | 1.00 | 0.95 | 0.99 | 0.78 | 0.07 |
| MAGPI-unopt | MEGE | 0.97 | 0.97 | 0.97 | 1.00 | 1.00 | 0.98 | 1.00 | 0.87 | 0.04 |
| MAGPI | MAGPI | 0.97 | 0.98 | 0.97 | 1.00 | 1.00 | 0.98 | 1.00 | 0.91 | 0.03 |

Overall, the consistently good performance across tube content, scan and rotation angle in the absence of phase estimation errors validates the Δ*χ* model (Eq 1) and demonstrates that the rotating-tube phantom itself did not introduce unexpected detrimental effects to the phase measurements. In previous work, QSM performance improved with higher isotropic spatial resolution and higher coverage^50, 55^. Here, slice thickness was 2.0mm for all scans, which may explain why performance did not drastically change with resolution. Additionally, Zhou et al^55^ and Karsa et al^50^ examined the entire QSM process, including inversion, which was not addressed here. Olsson et al^18^ used a phantom with tubes of Gd in comparable concentrations, though only one tube was used when varying angle with respect to B_0_ (five angles). That study used one method to estimate QSM^56-58^ over multiple spatial resolutions, volumes and inversion parameters. Similar to Karsa et al^50^, Olsson et al’s results improved with increasing resolution and volume coverage, and, similar to our results, Olsson et al observed errors in phase estimation using ^42^, compared to the theoretical result.

We introduced a wide range of variability to test repeatability and reproducibility of many data acquisition scenarios. While the performance of each method could be improved with additional ‘intervention’ and potentially adapting the acquisition parameters to the intended post-processing methods to be used later, our intent was to assess the ability of existing techniques in diverse imaging scenarios encountered in reality. *In vivo* imaging may present different sources of phase errors not included here (e.g., eddy currents, susceptibility-induced signal drops). The degree to which Steps 3 and 4 of QSM are sensitive to the errors introduced in Steps 1 and 2 requires further investigation. This phantom validation study allowed us to set a quantitative limit on the performance of various Step 1&2 methods. On-going work focuses on evaluating the performance of a subset of these methods, paired with Step 3 and 4 methods.

## 5 Conclusions

In this work, we used a rotating-tube phantom to explore sources of error in QSM data acquisition and phase estimation. To assess the robustness and repeatability of methods, we did not manually intervene. The two most impactful parameters on reproducibility of measurements were (a) acquisition protocol (e.g., single or multiple echoes) and (b) phase errors. The most repeatable and reproducible approaches were MAGPI and MAGPI-unopt, both methods based on the Maximum-Likelihood approach in phase estimation. For the remaining methods, performance varied greatly, even when systematically applied to the same underlying data from the same sequence or with the same method across different sequences. We intend to use the best-performing phase estimation methods from this experiment to analyze sources of error in background field removal (Step 3) and dipole inversion (Step 4).

## Appendix A Additional Experimental Details

## A.1 Frequency Referencing Methods

The Frequency Referencing method is attempting to remove the frequency offsets due to the rotating apparatus, and not the frequency offset due to susceptibility differences. After the effect of the rotating apparatus is removed, we fit the referenced frequency to the angle of rotation (Eq 1).

We call the process of removing 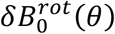 “frequency referencing”. We compute an estimate of 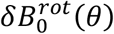 using the average frequency in a region outside the “Tube + Sphere” system. Our hypothesis is that this yields a good estimate of 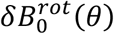 for three reasons: first, the region used for referencing (ROIs 1-13 in Fig. 2a) is static and not rotating with the apparatus. Second, the reference region is chosen far enough away from the Tube + Sphere system so that the “local susceptibility effects” (first term in Eq 2) do not affect the accuracy of the estimate of 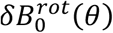. Finally, since the content in the reference region is homogenous material (water), the average field in each area improves the precision of this estimate.

In the table below, we include a few examples of 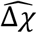 estimation using this model described in Eq 6.

**Table A.1.** 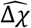 estimation using 1 mm resolution, TR=25 ms imaging data.

| Tube Method | 1 | 2 | 3 | 4 | 5 |
| --- | --- | --- | --- | --- | --- |
| $\Delta\chi_{theoretical}$ | 0.3362 | 0.3362 | 0.1681 | 0.1681 | 0.0804 |
| Slope $\widehat{\Delta\chi}$ | 0.2991 | 0.3177 | 0.1455 | 0.1868 | 0.0649 |
| MAGPI $\widehat{\Delta\chi}$ | 0.3157 | 0.2822 | 0.1496 | 0.1930 | 0.0709 |
| GBC $\widehat{\Delta\chi}$ | 0.3175 | 0.2985 | 0.1514 | 0.1761 | 0.0605 |
| MEDI.RG $\widehat{\Delta\chi}$ | 0.2970 | 0.3270 | 0.1422 | 0.1925 | 0.0595 |

Applying the susceptibility fit, after the frequency referencing step results in a good estimate of 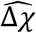, as demonstrated by the values in Table A.1.

## A.2 Determination of Reference Values

Susceptibility theory^59^ gives two approaches to determine *Δ*χ_th_ via Curie’s Law: first, using effective permeability, *μ*_*eff*_, based on experimentally determined values from the literature^60^ and second, using the permeability, μ, calculated from the spin orbitals. In the manuscript, we reported the values from the empirically-determined magnetic moment. For GdCl_3_, the theoretically-determined values from both methods were 0.168 ppm and 0.336 ppm, respectively. However, for CuSO_4_, the two theoretical values were 0.0804 ppm, determined using the highest reported Cu^2+^ moment, 2.17 Bohr magnetons, and 0.05112 ppm, determined using the spin orbital angular momentum quantum number, s=1/2, and the Landé g factor=2. We performed independent validation of the theoretical susceptibility values using NMR measurements.

Briefly, NMR susceptibility was determined from the chemical shift or peak separation between a H_2_O reference vial and the sample under test, each in capillaries placed in a standard 5 mm NMR tube. The NMR measurements were made at 14 T (600 MHz) and at 21.42 C (comparable to the MRI experimental temperature range 21.5 +/-0.5 C). The measured NMR susceptibility for 0.5 mM GdCl_3_ was 0.176 ppm and for 1.0 mM GdCl_3_ was 0.372 ppm; 5 % and 10 % greater than the theoretical values. The NMR-determined susceptibility for 3.2 mM CuSO_4_ in two separate experiments was 0.075 ppm and 0.0837 ppm. The NMR measurements for CuSO_4_ were 7 % less and 4 % greater than the *Δ*χ_th_ = 0.0804 ppm, and 47 % and 64 % greater than the *Δ*χ_th_ = 0.0511 ppm.

Copper has a low susceptibility value; CuSO_4_ is a more complex solution than GdCl_3_, and as a result, it is useful to test the lower limit of the susceptibility measurements. However, the moment in copper solutions is highly dependent on the local environment, which leads to the broader range of theoretical solutions. Based on the NMR measurements, the empirically-measurement moment is a better representation of the local environment in our experiment (including the CuSO_4_ solution, concentration and tube materials).

## Disclosures

Authors K.E.K, B.P.B, S.C., S.E.R., W.-T.W., J.A.B, and D.L.P. have nothing relevant to disclose. J.D. is the inventor of the MAGPI method and is the founder of MAGPI LLC.

## Acknowledgements

Authors W.-T.W., J.A.B., and D.L.P. would like to acknowledge funding from the Department of Defense in the Center for Neuroscience and Regenerative Medicine and the intramural research program of the National Institutes of Health.

## Code, Data, and Materials Availability

The authors will make the data and code available at a later date.

## References

1. Y. Wang, and T. Liu, “Quantitative susceptibility mapping (QSM): Decoding MRI data for a tissue magnetic biomarker,” Magnetic resonance in medicine : official journal of the Society of Magnetic Resonance in Medicine / Society of Magnetic Resonance in Medicine 73(1), 82–101 (2015) [10.1002/mrm.25358].

2. J. H. Jensen et al., “Magnetic field correlation as a measure of iron-generated magnetic field inhomogeneities in the brain,” Magnetic resonance in medicine : official journal of the Society of Magnetic Resonance in Medicine / Society of Magnetic Resonance in Medicine 61(2), 481–485 (2009) [10.1002/mrm.21823].

3. K. Shmueli et al., “Magnetic susceptibility mapping of brain tissue in vivo using MRI phase data,” Magnetic resonance in medicine : official journal of the Society of Magnetic Resonance in Medicine / Society of Magnetic Resonance in Medicine 62(6), 1510–1522 (2009) [10.1002/mrm.22135].

4. H. Tan et al., “Evaluation of iron content in human cerebral cavernous malformation using quantitative susceptibility mapping,” Invest Radiol 49(7), 498–504 (2014) [10.1097/RLI.0000000000000043].

5. M. Haacke et al., “Oxygen Saturation: Quantification,” in MRI: Basic Concepts and Clinical Applications M. Haacke, and J. Reichenbach, Eds., pp. 517–528, John Wiley & Sons, Inc (2011).

6. S. D. Sharma et al., “Quantitative susceptibility mapping in the abdomen as an imaging biomarker of hepatic iron overload,” Magnetic resonance in medicine : official journal of the Society of Magnetic Resonance in Medicine / Society of Magnetic Resonance in Medicine 74(3), 673–683 (2015) [10.1002/mrm.25448].

7. K. Deh et al., “Reproducibility of quantitative susceptibility mapping in the brain at two field strengths from two vendors,” Journal of magnetic resonance imaging : JMRI 42(6), 1592–1600 (2015) [10.1002/jmri.24943].

8. K. Deh et al., “Multicenter reproducibility of quantitative susceptibility mapping in a gadolinium phantom using MEDI+0 automatic zero referencing,” Magnetic resonance in medicine : official journal of the Society of Magnetic Resonance in Medicine / Society of Magnetic Resonance in Medicine 81(2), 1229–1236 (2019) [10.1002/mrm.27410].

9. X. Feng, A. Deistung, and J. R. Reichenbach, “Quantitative susceptibility mapping (QSM) and R2(*) in the human brain at 3T: Evaluation of intra-scanner repeatability,” Z Med Phys 28(1), 36–48 (2018) [10.1016/j.zemedi.2017.05.003].

10. T. Hinoda et al., “Quantitative Susceptibility Mapping at 3 T and 1.5 T: Evaluation of Consistency and Reproducibility,” Invest Radiol 50(8), 522–530 (2015) [10.1097/RLI.0000000000000159].

11. P. Y. Lin, T. C. Chao, and M. L. Wu, “Quantitative susceptibility mapping of human brain at 3T: a multisite reproducibility study,” AJNR. American journal of neuroradiology 36(3), 467–474 (2015) [10.3174/ajnr.A4137].

12. M. D. Santin et al., “Reproducibility of R2 * and quantitative susceptibility mapping (QSM) reconstruction methods in the basal ganglia of healthy subjects,” NMR in biomedicine 30(4), (2017) [10.1002/nbm.3491].

13. S. D. Robinson et al., “An illustrated comparison of processing methods for MR phase imaging and QSM: combining array coil signals and phase unwrapping,” NMR in biomedicine 30(4), (2017) [10.1002/nbm.3601].

14. R. Wang et al., “Stability of R2* and quantitative susceptibility mapping of the brain tissue in a large scale multi-center study,” Sci Rep 7(45261 (2017) [10.1038/srep45261].

15. M. Lancione et al., “Echo-time dependency of quantitative susceptibility mapping reproducibility at different magnetic field strengths,” NeuroImage 197(557-564 (2019) [10.1016/j.neuroimage.2019.05.004].

16. C. Langkammer et al., “Quantitative susceptibility mapping: Report from the 2016 reconstruction challenge,” Magnetic resonance in medicine : official journal of the Society of Magnetic Resonance in Medicine / Society of Magnetic Resonance in Medicine 79(3), 1661–1673 (2018) [10.1002/mrm.26830].

17. Z. Liu et al., “Preconditioned total field inversion (TFI) method for quantitative susceptibility mapping,” Magnetic resonance in medicine : official journal of the Society of Magnetic Resonance in Medicine / Society of Magnetic Resonance in Medicine 78(1), 303–315 (2017) [10.1002/mrm.26331].

18. E. Olsson, R. Wirestam, and E. Lind, “MRI-Based Quantification of Magnetic Susceptibility in Gel Phantoms: Assessment of Measurement and Calculation Accuracy,” Radiol Res Pract 2018(6709525 (2018) [10.1155/2018/6709525].

19. V. Fortier, and I. R. Levesque, “Phase processing for quantitative susceptibility mapping of regions with large susceptibility and lack of signal,” Magnetic resonance in medicine : official journal of the Society of Magnetic Resonance in Medicine / Society of Magnetic Resonance in Medicine 79(6), 3103–3113 (2018) [10.1002/mrm.26989].

20. H. Erdevig et al., “Accuracy of magnetic resonance based susceptibility measurements,” AIP Advances 7(5), 056718 (2017)

21. Y. C. Cheng et al., “Quantifying effective magnetic moments of narrow cylindrical objects in MRI,” Physics in medicine and biology 54(22), 7025–7044 (2009) [10.1088/0031-9155/54/22/018].

22. C. Liu et al., “Susceptibility-weighted imaging and quantitative susceptibility mapping in the brain,” Journal of magnetic resonance imaging : JMRI 42(1), 23–41 (2015) [10.1002/jmri.24768].

23. S. Liu et al., “Susceptibility-weighted imaging: current status and future directions,” NMR in biomedicine 30(4), (2017) [10.1002/nbm.3552].

24. U. Katscher, T. Voigt, and C. Findeklee, “Electrical conductivity imaging using magnetic resonance tomography,” Conf Proc IEEE Eng Med Biol Soc 2009(3162–3164 (2009) [10.1109/IEMBS.2009.5334031].

25. V. Rieke, and K. Butts Pauly, “MR thermometry,” Journal of magnetic resonance imaging : JMRI 27(2), 376–390 (2008) [10.1002/jmri.21265].

26. S. Ley et al., “Value of MR phase-contrast flow measurements for functional assessment of pulmonary arterial hypertension,” Eur Radiol 17(7), 1892–1897 (2007) [10.1007/s00330-006-0559-9].

27. R. Muthupillai et al., “Magnetic resonance elastography by direct visualization of propagating acoustic strain waves,” Science 269(5232), 1854–1857 (1995) [10.1126/science.7569924].

28. E. M. Haacke et al., “Quantitative susceptibility mapping: current status and future directions,” Magnetic resonance imaging 33(1), 1–25 (2015) [10.1016/j.mri.2014.09.004].

29. P. Sati et al., “Rapid, high-resolution, whole-brain, susceptibility-based MRI of multiple sclerosis,” Mult Scler 20(11), 1464–1470 (2014) [10.1177/1352458514525868].

30. W. Feng, J. Neelavalli, and E. M. Haacke, “Catalytic multiecho phase unwrapping scheme (CAMPUS) in multiecho gradient echo imaging: removing phase wraps on a voxel-by-voxel basis,” Magnetic resonance in medicine : official journal of the Society of Magnetic Resonance in Medicine / Society of Magnetic Resonance in Medicine 70(1), 117–126 (2013) [10.1002/mrm.24457].

31. G. Helms, and P. Dechent, “Increased SNR and reduced distortions by averaging multiple gradient echo signals in 3D FLASH imaging of the human brain at 3T,” Journal of magnetic resonance imaging : JMRI 29(1), 198–204 (2009) [10.1002/jmri.21629].

32. J. Dagher, and K. Nael, “MAGPI: A framework for maximum likelihood MR phase imaging using multiple receive coils,” Magnetic resonance in medicine : official journal of the Society of Magnetic Resonance in Medicine / Society of Magnetic Resonance in Medicine 75(3), 1218–1231 (2016) [10.1002/mrm.25756].

33. J. Dagher, and K. Nael, “MR phase imaging with bipolar acquisition,” NMR in biomedicine 30(4), (2017) [10.1002/nbm.3523].

34. A. M. Halefoglu, and D. M. Yousem, “Susceptibility weighted imaging: Clinical applications and future directions,” World J Radiol 10(4), 30–45 (2018) [10.4329/wjr.v10.i4.30].

35. C. J. Bakker, H. de Leeuw, and P. R. Seevinck, “Selective depiction of susceptibility transitions using Laplace-filtered phase maps,” Magnetic resonance imaging 30(5), 601–609 (2012) [10.1016/j.mri.2011.12.023].

36. N. K. Chen, and A. M. Wyrwicz, “Correction for EPI distortions using multi-echo gradient-echo imaging,” Magnetic resonance in medicine : official journal of the Society of Magnetic Resonance in Medicine / Society of Magnetic Resonance in Medicine 41(6), 1206–1213 (1999) [10.1002/(sici)1522-2594(199906)41:6<1206::aid-mrm17>3.0.co;2-l].

37. R. M. Goldstein, H. A. Zebker, and C. L. Werner, “Satellite Radar Interferometry - Two-Dimensional Phase Unwrapping,” Radio Sci 23(4), 713–720 (1988) [DOI 10.1029/RS023i004p00713].

38. K. Lu, T. T. Liu, and M. Bydder, “Optimal phase difference reconstruction: comparison of two methods,” Magnetic resonance imaging 26(1), 142–145 (2008) [10.1016/j.mri.2007.04.015].

39. M. A. Schofield, and Y. Zhu, “Fast phase unwrapping algorithm for interferometric applications,” Opt Lett 28(14), 1194–1196 (2003) [10.1364/ol.28.001194].

40. Y. Wang, “Cornell MEDI Toolbox,” (2019).

41. S. Witoszynskyj et al., “Phase unwrapping of MR images using Phi UN--a fast and robust region growing algorithm,” Med Image Anal 13(2), 257–268 (2009) [10.1016/j.media.2008.10.004].

42. R. Cusack, and N. Papadakis, “New robust 3-D phase unwrapping algorithms: application to magnetic field mapping and undistorting echoplanar images,” NeuroImage 16(3 Pt 1), 754–764 (2002)

43. B. Wu et al., “Fast and tissue-optimized mapping of magnetic susceptibility and T2* with multi-echo and multi-shot spirals,” NeuroImage 59(1), 297–305 (2012) [10.1016/j.neuroimage.2011.07.019].

44. J. L. R. Andersson et al., “Susceptibility-induced distortion that varies due to motion: Correction in diffusion MR without acquiring additional data,” NeuroImage 171(277–295 (2018) [10.1016/j.neuroimage.2017.12.040].

45. J. A. Lundman et al., “Patient-induced susceptibility effects simulation in magnetic resonance imaging,” Physics and Imaging in Radiation Oncology 1(41–45 (2017) [j.phro.2017.02.004].

46. P. B. Roemer et al., “The NMR phased array,” Magnetic resonance in medicine : official journal of the Society of Magnetic Resonance in Medicine / Society of Magnetic Resonance in Medicine 16(2), 192–225 (1990) [10.1002/mrm.1910160203].

47. B. Gruber et al., “RF coils: A practical guide for nonphysicists,” Journal of magnetic resonance imaging : JMRI (2018) [10.1002/jmri.26187].

48. S. C. Chu et al., “Bulk magnetic susceptibility shifts in NMR studies of compartmentalized samples: use of paramagnetic reagents,” Magnetic Resonance in Medicine 13(2), 239–262 (1990)

49. C. Y. Hsieh et al., “An improved method for susceptibility and radius quantification of cylindrical objects from MRI,” Magnetic resonance imaging 33(4), 420–436 (2015) [10.1016/j.mri.2015.01.004].

50. A. Karsa, S. Punwani, and K. Shmueli, “The effect of low resolution and coverage on the accuracy of susceptibility mapping,” Magnetic resonance in medicine : official journal of the Society of Magnetic Resonance in Medicine / Society of Magnetic Resonance in Medicine 81(3), 1833–1848 (2019) [10.1002/mrm.27542].

51. M. J. Cronin et al., “Exploring the origins of echo-time-dependent quantitative susceptibility mapping (QSM) measurements in healthy tissue and cerebral microbleeds,” NeuroImage 149(98–113 (2017) [10.1016/j.neuroimage.2017.01.053].

52. Y. Kanazawa et al., “Appropriate echo time selection for quantitative susceptibility mapping,” Radiol Phys Technol 12(2), 185–193 (2019) [10.1007/s12194-019-00513-x].

53. W. Zheng et al., “Measuring iron in the brain using quantitative susceptibility mapping and X-ray fluorescence imaging,” NeuroImage 78(68–74 (2013) [10.1016/j.neuroimage.2013.04.022].

54. M. C. Hsieh et al., “Quantitative Susceptibility Mapping-Based Microscopy of Magnetic Resonance Venography (QSM-mMRV) for In Vivo Morphologically and Functionally Assessing Cerebromicrovasculature in Rat Stroke Model,” PLoS One 11(3), e0149602 (2016) [10.1371/journal.pone.0149602].

55. D. Zhou et al., “Susceptibility underestimation in a high-susceptibility phantom: Dependence on imaging resolution, magnitude contrast, and other parameters,” Magnetic resonance in medicine : official journal of the Society of Magnetic Resonance in Medicine / Society of Magnetic Resonance in Medicine 78(3), 1080–1086 (2017) [10.1002/mrm.26475].

56. L. de Rochefort et al., “Quantitative susceptibility map reconstruction from MR phase data using bayesian regularization: validation and application to brain imaging,” Magnetic resonance in medicine : official journal of the Society of Magnetic Resonance in Medicine / Society of Magnetic Resonance in Medicine 63(1), 194–206 (2010) [10.1002/mrm.22187].

57. J. Liu et al., “Morphology enabled dipole inversion for quantitative susceptibility mapping using structural consistency between the magnitude image and the susceptibility map,” NeuroImage 59(3), 2560–2568 (2012) [10.1016/j.neuroimage.2011.08.082].

58. T. Liu et al., “Nonlinear formulation of the magnetic field to source relationship for robust quantitative susceptibility mapping,” Magnetic resonance in medicine : official journal of the Society of Magnetic Resonance in Medicine / Society of Magnetic Resonance in Medicine 69(2), 467–476 (2013) [10.1002/mrm.24272].

59. R. Carlin, “Magnetochemistry,” Springer, New York, NY (1986).

60. G. A. Bain, and J. F. Berry, “Diamagnetic corrections and Pascal’s constants,” J Chem Educ 85(4), 532–536 (2008) [DOI 10.1021/ed085p532].

